# Recognition of non-self is necessary to activate *Drosophila*’s immune response against an insect parasite

**DOI:** 10.1101/2022.06.28.497890

**Authors:** Alexandre B. Leitão, Ramesh Arunkumar, Jonathan P. Day, Nancy Hanna, Aarathi Devi, Matthew P. Hayes, Francis Jiggins

**Affiliations:** Department of Genetics, University of Cambridge, Cambridge, United Kingdom; Champalimaud Foundation, Portugal; Royal College of Surgeons in Ireland, Ireland; Department of Zoology, University of Cambridge, Cambridge, United Kingdom

## Abstract

Innate immune responses can be activated by pathogen-associated molecular patterns (PAMPs) or danger signals released by damaged tissues. As PAMPs are typically conserved across broad groups of pathogens but absent from the host, it is unclear whether they allow hosts to recognize parasites that are phylogenetically related to themselves, such as parasitoid wasps infecting insects. Parasitoids must penetrate the cuticle of *Drosophila* larvae to inject their eggs. In line with previous results, we find that the danger signal of wounding triggers the differentiation of specialized immune cells called lamellocytes. However, using oil droplets to mimic infection by a parasitoid wasp egg, we find that the activation of melanization response that kills parasitoids also requires exposure to a parasitoid wasp molecule that acts as a PAMP. The unidentified factor enhances the transcriptional response in hemocytes and induces a specific response in the fat body that includes *Tep1*, which is essential for efficient melanization. We conclude that a combination of danger signals and PAMPs are required activate *Drosophila*’s immune response against parasitic insects.

## Introduction

Organisms must be able to reliably detect when they are infected in order to mount an appropriate immune response, and this frequently relies on the recognition of non-self. In the case of innate immune systems, pattern recognition receptors (PRRs) detect pathogen-associated molecular patterns (PAMPs). These are typically molecules such as flagellin or peptidoglycan that are absent from the host but highly conserved across a broad class of pathogens [1]. An alternative way to sense infection is to detect danger signals such as cell damage. Here, damaged cells release damage-associated molecular patterns (DAMPs), which bind host receptors and trigger the immune response [2]. Finally, pathogens may be detected because of ‘missing self’—they lack some factor found on host cells that inhibits immune activation [3].

Sometimes, immune responses must be mounted against parasites that are closely related to the host. For example, plants can be infected by other plants, insects by other insects, and some mammals are even infected by transmissible cancer cells derived from their own species [4]. In one of the few cases that have been characterized, tomato plants have evolved a PRR to recognize a PAMP produced by the pathogenic plant *Cuscuta reflexa* [5,6]. However, it is unclear whether this will be more broadly true. In many cases, it may be difficult to recognize PAMPs as there will be fewer conserved differences between closely related pairs of hosts and pathogens. This problem may be exacerbated as there is selection on the pathogen to escape recognition by losing their PAMPs, and this may be easier to evolve if you are already similar to your host. If this is the case, recognition may rely on danger signals or detecting missing-self.

This problem is especially acute for insects, as their most important parasites are frequently other insects [7]. Many parasitoid wasps infect their insect hosts by injecting eggs into their haemocoel. Typically, infection by parasitoids leads to activation of an immune response that involves the formation of a cellular capsule around the parasitoid egg that later becomes melanized [8–10]. The cellular immune response to parasitoid wasps in *Drosophila melanogaster* involves the differentiation of a hemocyte type rarely found in healthy larvae, the lamellocyte [8]. These form the outer-layer of the cellular capsule around the parasitoid egg, which is melanized when pro-phenoloxidase 2 and 3 (PPO2 and PPO3) are activated in crystal cells and lamellocytes respectively [11].

The recognition of parasitoid infections relies in part on danger signals. Lamellocyte differentiation can be triggered by sterile wounding of the larval cuticle [12], which presumably mimics a parasitoid piercing the cuticle with its ovipositor. Furthermore, in many insects, introducing inert objects into the haemocoel leads to a cellular encapsulation response [13], and in species like *D. yakuba*, this is accompanied by the object being melanized [14,15]. However, in other cases these danger signals only lead to an incomplete immune response, as a cellular capsule forms but is not melanized [16], or, in species like *D. melanogaster*, there is only a low level of melanization [15,17]. Interestingly, *D. melanogaster* larvae that have been parasitized by the wasp *Asobara tabida* are more likely to strongly melanize inert objects [17]. This suggests that a PAMP injected by the parasitoid might activate the melanization response. Here, we examine how the combination of danger signals and PAMPs activate different components of the immune response of *D. melanogaster* to parasitoid wasps.

## Results

### Danger signals induce immune cell differentiation

Parasitoid wasp attack induces the rapid differentiation of blood cells called lamellocytes, which encapsulate and melanize the wasp. It has previously been reported that sterile wounding of larval cuticle induces lamellocyte differentiation [12]. In line with this, we found that injecting a droplet of paraffin oil induced lamellocyte differentiation (Figure 1A; main effect of treatment: *F* = 36.187, d.f = 2, 38, *p* = 1.59 × 10^−9^; oil vs. control *t* = 5.298, d.f. = 38, *p*<1.57×10^−5^).

**Figure 1.**
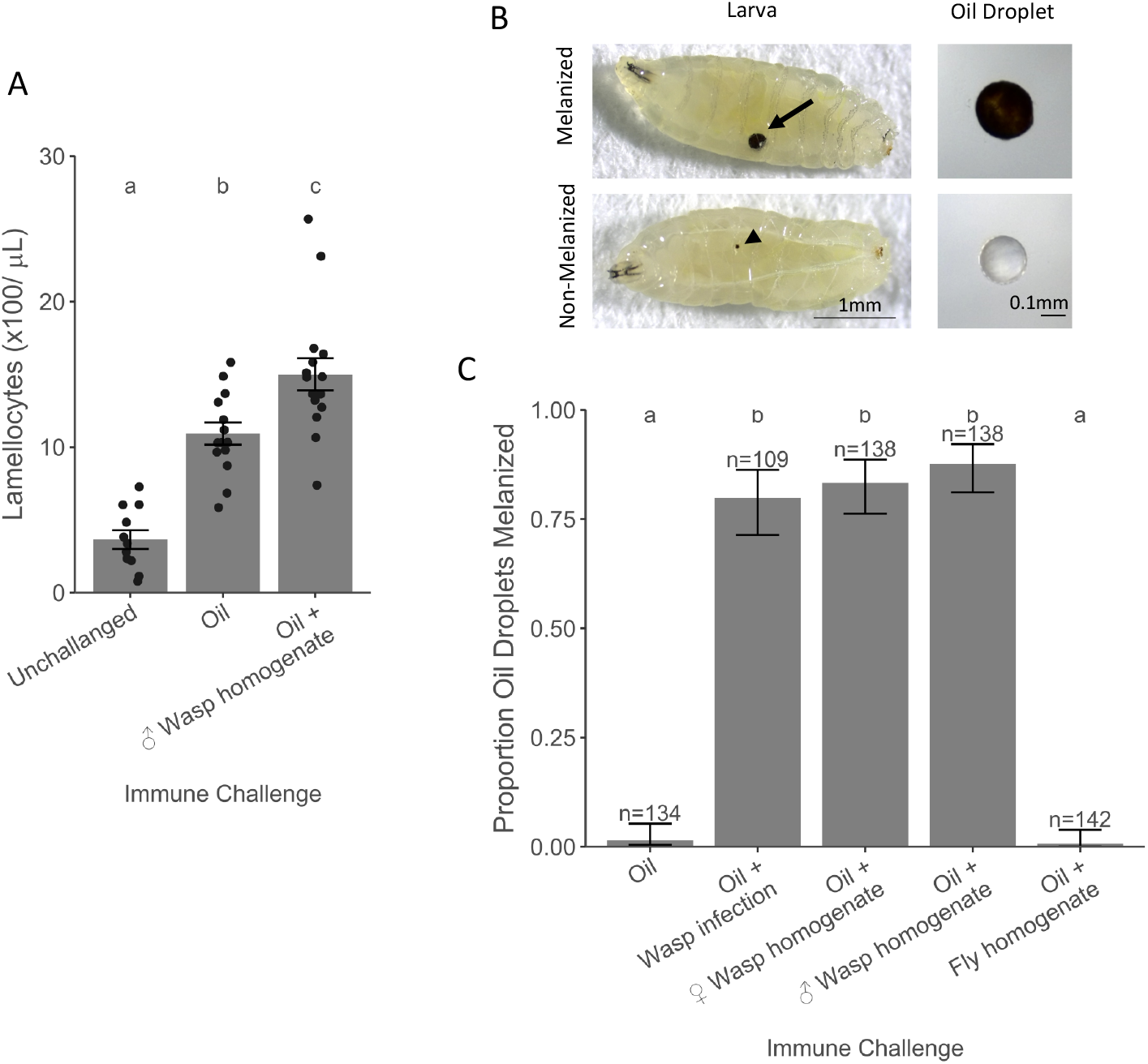
The effect of parasitoid wasp exposure on the melanization of oil droplets. (A) Concentration of lamellocytes in the hemolymph of unchallenged larvae and larvae 48h post injection with oil or oil + wasp homogenate. The data points are independent measurements on hemolymph pooled from 8-10 larvae. (B) Oil droplets injected into larvae are either melanized (arrow) or not. Melanization of cuticle resulting from injection wounding is often visible (arrow head). (C) Proportion of larvae with melanized oil droplets 48h after different immune challenges. Different letters represent treatments with statistically significant differences (Tukey’s Honest Significant Difference Test, *p*<0.01).

To examine the role of PAMPs in this response, we homogenized adult male wasps in the paraffin oil before injecting the fly larvae. Therefore, this treatment combines both the wounding from injection and exposure to PAMPs. We found that the addition of wasp homogenate led to a larger number of lamellocytes being produced (Figure 1A; oil vs. wasp homogenate *t* = 3.26, d.f. = 38, *p*=0.007). This suggests that danger signals resulting from the wound created by the female ovipositor are the primary factor triggering lamellocyte differentiation, but the response is amplified by recognition of non-self from wasps.

### Parasite molecules activate the melanization response

The final step of the immune response against parasitoid wasps is the melanization of the wasp egg. To test if this requires the recognition of non-self, we examined whether paraffin oil droplets were melanized 48h after injection (Figure 1B). Wounding alone was not sufficient to activate this response, as paraffin oil by itself did not induce a strong melanization reaction (Figure1C). However, if larvae were previously infected by a low virulence *L. boulardi* strain (G486), the melanization of the oil droplet increases (Figure1C; main effect of treatment: *X*^*2*^ = 577.39, d.f = 4, *p* < 2 × 10^−16^; oil vs. oil + wasp infection: z = 7.612, *p* = 2 × 69^-13^). Therefore, the presence of the parasite is required to trigger this immune response.

To test whether parasitoid wasp PAMPs are responsible for the activation of the immune response, we injected flies with paraffin oil containing homogenized female wasps. This led to a robust melanization response that was indistinguishable from that seen when the flies had been parasitized (Figure 1C; female wasp homogenate vs oil: *z*=7.967, *p* = 1 × 55^-14^). Furthermore, this reaction is not due to the presence of eggs or venoms in the female wasp homogenates, as male wasp homogenates induced a similar response (Figure 1C; male homogenate vs female homogenate: z=1.051, *p*=1).

The immune response to our crude homogenate of parasitoid wasp could be a specific response to PAMPs in the parasitoid tissue or a general response to injecting damaged cells, which are known to release DAMPs that activate the innate immune system [18]. To distinguish these hypotheses, we injected larvae with paraffin oil containing *D. melanogaster* homogenate. This did not induce melanization (Figure 1C; oil vs fly homogenate: z=-0.634, *p*=1). Together, these results indicate that the *D. melanogaster* immune system recognizes non-self molecules in the parasitoid wasp to activate the melanization response.

To determine if the fly immune system recognizes wasp proteins, we treated the wasp homogenate with proteinase K, a broad range serine protease [19]. This did not reduce the proportion of melanized oil droplets (Figure S1, main effect of treatment: *X*^*2*^ = 102.24, d.f = 4, *p* < 2 × 10^−16^; No autoclave vs. No autoclave + Proteinase K: z = 1.267, *p* =1). Serendipitously, we tested samples where wasps were autoclaved before homogenization. In itself this has no effect on melanization rates (Autoclave vs. No autoclave: z = 1.254, *p* =1). However, when the autoclaved homogenate is treated with proteinase K, the number of melanized oil droplets is significantly reduced (Autoclave vs. Autoclave + Proteinase K: z = 5.41, *p* =6.3 × 10^−7^). This result suggests that one or more proteins in the wasp body act as PAMPs to activate the fly immune system.

### PAMPs activate the humoral immune response

In addition to the cellular immune response, the melanization of the capsule formed around wasp eggs relies on a humoral immune response involving the secretion of molecules from the fat body [20]. To understand the effects of danger signals and wasp PAMPs on this response, we sequenced RNA extracted from fat body 24h post injection of paraffin oil, paraffin oil with wasp homogenate and from non-injured controls (unchallenged). After aligning the RNA-seq reads to the *D. melanogaster* genome, the number of uniquely mapped exonic reads ranged from 3,288,236 to 15,962,793 (Table 1B).

The injection of paraffin oil alone did not lead to the significant upregulation of any genes at 24h, but the addition of wasp homogenate upregulated 29 genes (Figure 2A-B). Ten of these encode serine proteases with a trypsin domain (Table S2), a class of proteins known to be involved in the melanization cascade and Toll pathway [20]. Other upregulated genes encode immunity-related molecules, including Toll, thioester-containing proteins (TEPs) and a fibrinogen.

**Figure 2.**
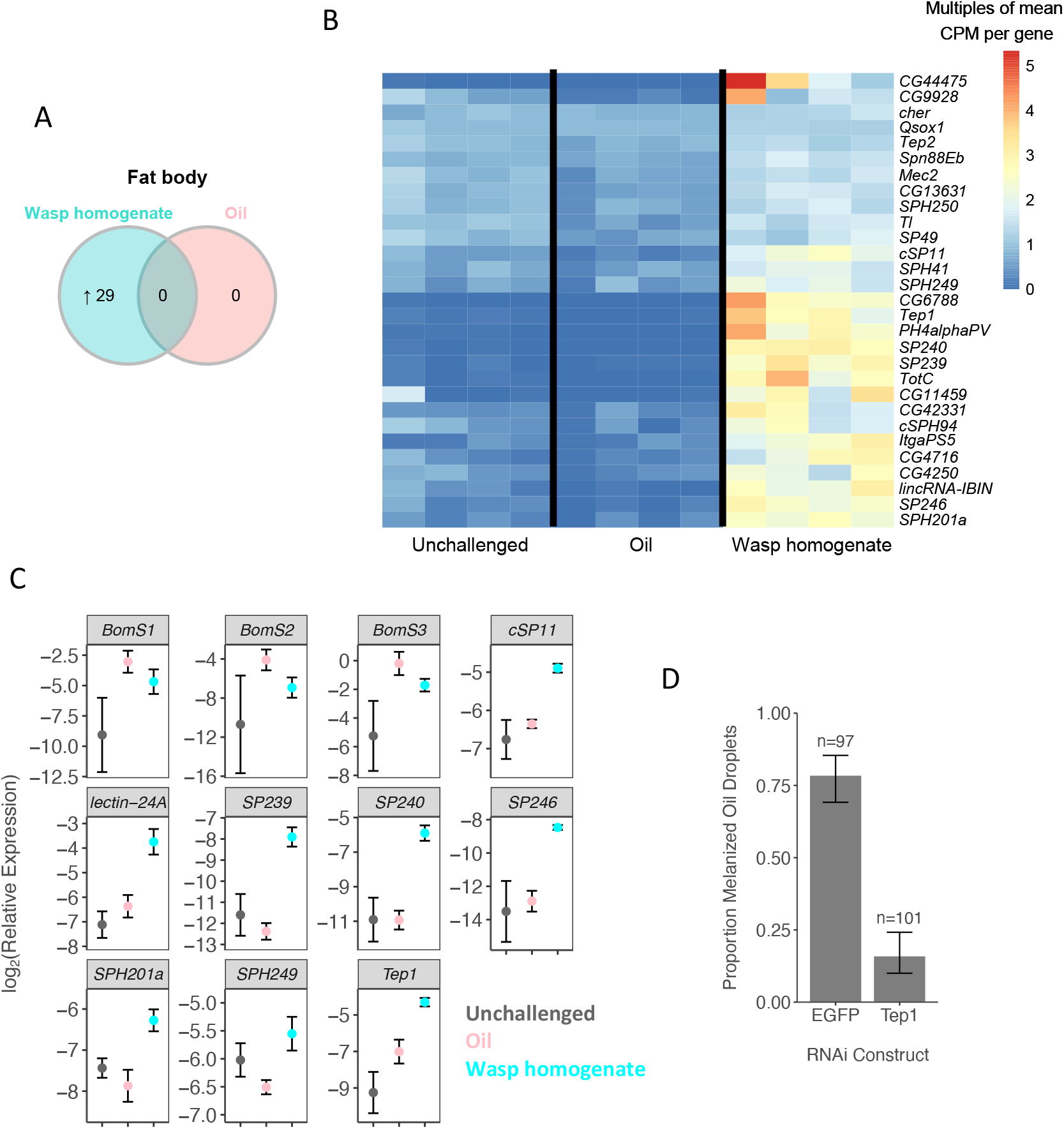
Transcriptional response of fat body to wasp exposure. Larvae were injected with oil, wasp + oil or unchallenged. RNA from the fat body was sequenced 24h post treatment. (A) The number of genes with significantly changes in expression compared with unchallenged conditions. (B) The expression of 29 genes with significantly changes in expression after immune challenge. (C) The rate of melanization of wasp + oil droplets in transgenic flies that express constructs to knock down the expression of *Tep1* or eGFP (control).

To confirm the specific response to wasp homogenate, we chose a subset of the genes and analysed gene expression with qPCR. This revealed that wounding and the wasp PAMP elicit distinct humoral immune responses. The injection of paraffin oil alone was sufficient to upregulate Bomanin genes, which encode short peptides that play a role in killing bacterial pathogens [21] (Figure 2C; Unchallenged vs Oil: *p=*0.04, *p*=0.05 and *p*=0.10). However, other genes including secreted serine proteases, a lectin and a Tep were specifically upregulated in the presence of the wasp homogenate (Figure 2C). This suggests that a humoral immune response against bacteria can to triggered by wounding, but PAMPs are required for the specific anti-parasitoid response.

Genes upregulated upon wasp homogenate injection include *Tep1* and *Tep2* (Figure 2B-C), which encode secreted complement-like proteins. TEPs are involved in resistance against bacteria, fungi and parasitoid wasps [22,23], and in the mosquito *Anopheles gambiae* they act as an opsinin, binding to the ookinetes of *Plasmodium* eggs [24]. We therefore tested the role of *Tep1* in the melanization process. Knocking down *Tep1* expression with RNAi reduced the ability of larvae to melanize the oil droplet prepared with wasp homogenate (Figure 2D, main effect of treatment: *X*^*2*^ = 16.367, d.f = 1, *p* = 4.525 × 10^−5^). Therefore, the wasp PAMPs upregulate genes required for the immune response that kills the parasite.

### Parasitoid wasp PAMPs amplify the transcriptional response of immune cells to danger signals

To understand the role of PAMPs in the cellular immune response we repeated the RNA-seq experiment on hemocytes. The number of uniquely mapped exonic reads ranged from 2,520,358 to 22,036,188 (Table 1B). There is a much broader transcriptional change in hemocytes than in the fat body (Figure 3A), with 3,887 genes being differentially expressed after larvae were injected with wasp homogenate (Figure 3A, Figure S2). The genes that were significantly differentially expressed when larvae were injected with mineral oil alone were largely a subset of these genes (Figure 3A). The wasp homogenate and mineral oil injections are largely causing the same genes to change in expression, but the magnitude of the transcriptional response is greater in the presence of the PAMP (Figure 3B). Therefore, the PAMP amplifies the response to a danger signal.

**Figure 3.**
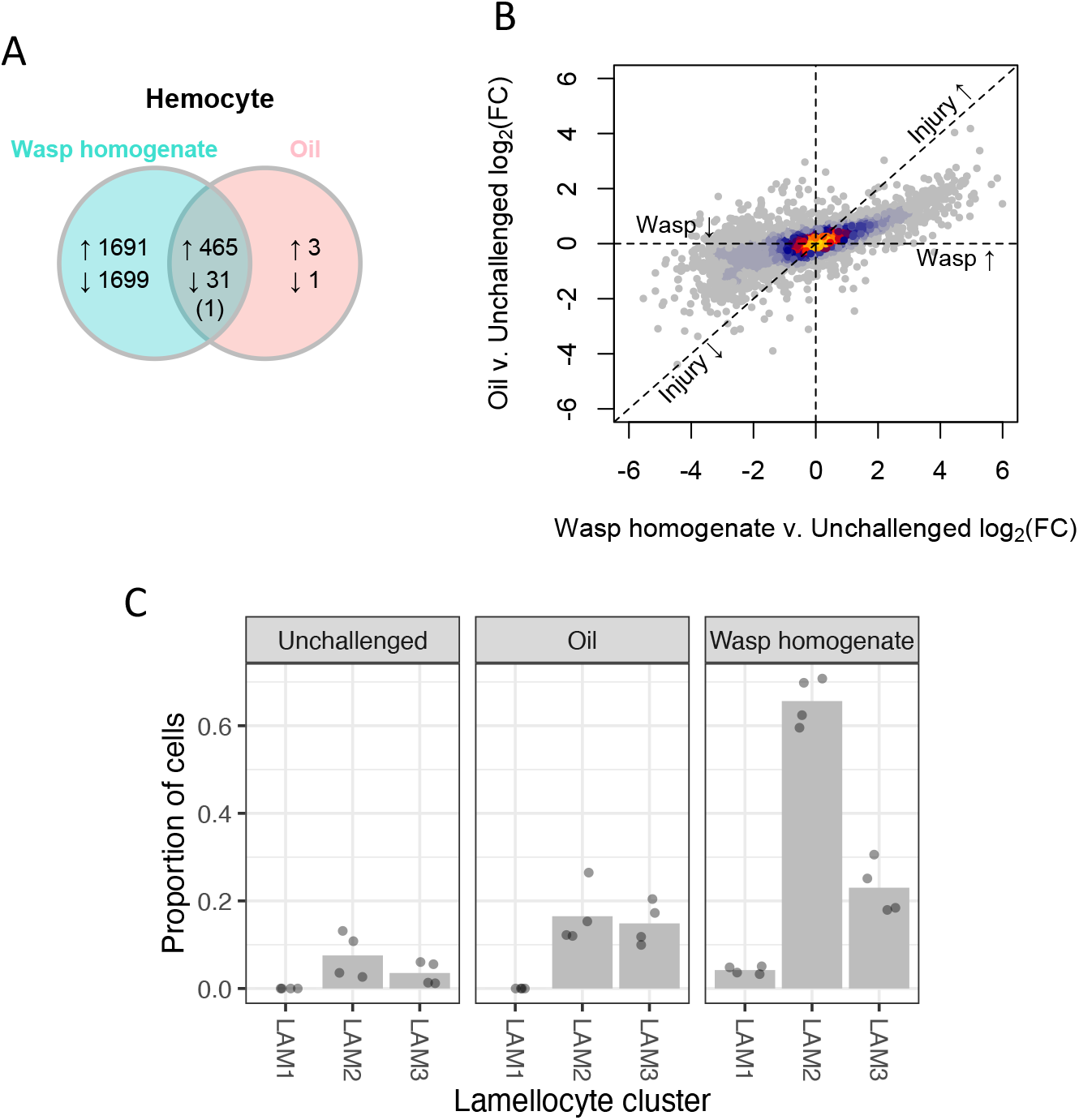
Transcriptional response of hemocytes to wasp exposure. Larvae were injected with oil, wasp + oil or unchallenged. RNA from the hemocytes was sequenced 24h post treatment. (A) The number of genes with significantly changes in expression compared with unchallenged conditions. (B) Changes in gene expression induced by injection of wasp homogenate (x-axis) and by injection of oil (y-axis). Because both treatments cause injury, genes solely regulated by injury will be close to the dashed diagonal 1:1 line. Genes specifically activated by wasp PAMPs will be on the x-axis. Relative expression is represented as log_2_(Fold Change). (C) Inferred proportion of immature (LAM1 and LAM2) and mature (LAM3) lamellocytes estimated from the RNA-seq data using digital cytometry. Each point is an independent sample and the bars are the mean.

The massive transcriptional response of hemocytes may reflect the differentiation of lamellocytes, which are rare in homeostasis but increase after wasp infection. Genes upregulated by wasp homogenate were enriched for biological adhesion and cytoskeleton organization, which may be related to the role of lamellocytes in capsule formation and the changes in cell morphology that occur as these cells differentiate (Figure S3A). The upregulated genes were also enriched for endocytosis, macroautophagy and other immune functions (Table S3) [25]. In contrast, genes downregulated by wasp homogenate were enriched for extracellular structure organization, a housekeeping function of plasmatocytes (Figure S3A).

To test whether these transcriptional changes were linked to the differentiation of lamellocytes, we compared our data to previous results we have generated using single-cell RNA sequencing (scRNA-seq) [26]. We found that genes that were highly expressed in lamellocytes were upregulated by injecting wasp homogenate and vice-versa for downregulated genes (Figure S3B). To investigate this further, we estimated the abundance of different hemocyte types using digital cytometry [27]. This is a statistical technique that estimate cell proportions in ‘bulk’ RNA-seq data using the single cell expression profile [26] as a reference. We estimated that there was a moderate increase in the proportion of lamellocytes following the injection of an oil droplet (Figure 3C). However, the injection of wasp homogenate led to the differentiation of mature lamellocytes (LAM3) together with large numbers of immature lamellocytes (Figure 3C).

### *Drosophila melanogaster* has evolved to recognize parasitoid-specific molecules

To test if *Drosophila* larvae have evolved to recognize parasitoid-specific molecules, we injected fly larvae with homogenates from prepared from 44 insect species and examined whether they activated the melanization response to oil droplets. The two parasitoid species resulted in the two highest melanization rates of all 44 species, and this parasitoid-specific response was significant after correcting for the phylogenetic relatedness of the 44 species (Figure 4; phylogenetic mixed model: *p*=0.006). Therefore, the *Drosophila* immune system appears to have evolved a specific mechanism to recognize parasitoids.

**Figure 4.**
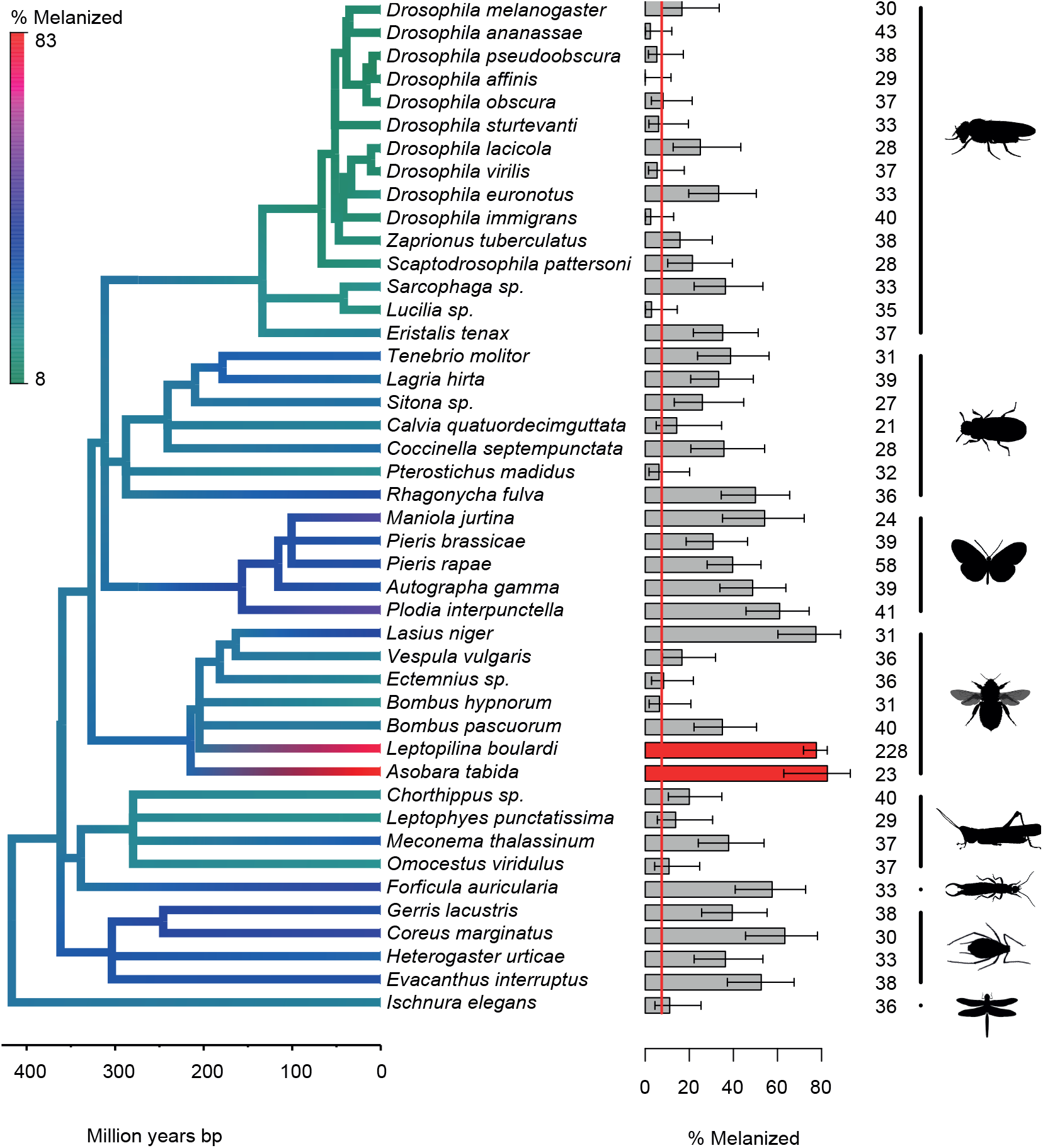
The effect of injecting homogenates of different insects on the melanization of oil droplets. The bar chart shows the proportion of oil droplets that were melanized with 95% binomial confidence intervals. The red line is the rate at which oil droplets were melanized without any insect homogenate (*N*=241). The red bars are parasitoids of *D. melanogaster*. Sample sizes are given beside the bars. The silhouettes represent different insect orders (from top: Diptera, Coleoptera, Lepidoptera, Hymenoptera, Orthoptera, Dermaptera, Hemiptera, Odonata).

Despite this parasitoid-specific response, homogenates of many other insect species cause some level of immune activation (Figure 4). It is striking that species closely related to *D. melanogaster* do not activate the melanization response (Figure 4; 95% confidence intervals overlap the dashed line basal response). Therefore, it appears that many insect species contain factors that cause some activation of the melanization response, but *Drosophila* immunity is not activated by self.

## Discussion

Immune responses are multistep processes that require several levels of regulation. Here, we describe how the immune response of *D. melanogaster* against parasitoid wasps is regulated by two modes of immune activation. In line with previous results [12], immune challenge with an inert object induces the differentiation of lamellocytes. The signal required for this is likely a damage associated molecular pattern (DAMP) produced during wounding [28]. However, the final step of the encapsulation response—melanization—only occurs when wasp molecules are present. This suggests that this part of the encapsulation response is activated by a pathogen associated molecular pattern (PAMP). This PAMP has different effects on the two main immune tissues of *Drosophila*. In hemocytes, it enhances the differentiation of lamellocytes, amplifying the effects wounding. In the fat body it triggers the upregulation of a small number of humoral immune genes whose expression is not affected by wounding alone. This includes the secreted complement-like molecule TEP1, which we show is required for the melanization response.

The reason why this immune response relies on both PAMPs and DAMPs is unknown. However, the PAMP may be used to adjust the immune response to target this specific parasite, as wounding may occur for many other reasons. Alternatively, it may allow the response to be modulated according to the levels of infection (although this may be less relevant to parasitoids than other forms of infection). This regulation step may be important for the fly to minimize costs associated with the activation of the melanization cascade. Artificial activation of the melanization response is detrimental for host tissue physiology [29], while the production of lamellocytes, that is enhanced by the PAMP, is thought to be energetically costly for the fly [30]. Without this step of immune regulation, any cuticular wound would result in activation of the melanization cascade and incur associated costs.

Homogenates prepared with other insect species can also result in melanization, but parasitoid wasps induce the highest response. This variation in the phenotype may be the result of recognition of different molecules or different quantities of the same molecule in different species. It is worth noticing that homogenates prepared with other *Drosophila* species induce a very poor response. This is in contrast with a previous report that shows encapsulation and melanization of fat bodies from heterospecific tissue transplants when the donor is a *Drosophila* species outside the melanogaster group [31]. In this case, host larvae are mutants in an unidentified gene that causes the differentiation of lamellocytes in homeostasis, possibly explaining the discrepancy with the results presented here.

All major macromolecules, proteins, carbohydrates, nucleic acids, and lipids can act as PAMPs [1], and the crude wasp homogenate used in our experiments includes all these molecules. However, we found that treatment with proteinase K reduces the activity of the PAMP, suggesting that the active molecule may be a protein or glycoprotein. Unexpectedly, the proteinase K was only effective if the wasps were autoclaved prior to homogenization. This suggests that the wasp protein may be resistant to proteinase K digestion. Proteinase K resistant proteins are rare [32] but they include some eukaryotic proteins [33]. The high temperature treatment may have modified the configuration of the wasp PAMP and made it accessible to proteinase K digestion.

In conclusion, we show that *Drosophila*’s immune system can recognize the presence of wasp parasites and uses this recognition to modulate the cellular and humoral responses that are initiated by injury. To our knowledge, this is the first report of an animal immune system recognizing a PAMP from a parasite that is closely related to itself.

## Material and methods

### *D. melanogaster* and *L. boulardi* maintenance

For all experiments except RNAi we used an outbred *D. melanogaster* population that was established from 372 isofemales caught in Cambridgeshire in October 2017. Population size was maintained over 500 flies per generation and had over 10 generations of laboratory adaptation before the start of experiments. For experimental procedures, flies were allowed to lay eggs overnight on agar plates covered with yeast (*Saccharomyces cerevisiae* – Sigma YSC2). Eggs were washed from the agar plate with PBS and transferred into 1.5ml microcentrifuge tubes. 13μl of eggs and PBS (∼150 eggs) were transferred onto 50mm cornmeal food plates. These were incubated for 72 hours before experiments. Developing and adult *D. melanogaster* were maintained at 25°C, 70% relative humidity in an 8h-16h dark-light cycle.

*L. boulardi* was maintained by allowing females to infect 1^st^ instar larvae of the outbred population and incubating them at 25°C. Adult wasps were collected 24 days after infection and maintained at room temperature with a drop of honey for up a maximum of 5 days before infections. To infect *D. melanogaster* larvae, 3 *L. boulardi* females were allowed to infect larvae on the cornmeal food plates for 3 hours.

### RNAi

The following *D*.*melanogaster* stocks were obtained from the Bloomington Drosophila Stock Center: UAS-Tep1^dsRNA^ (BL#: 32856), UAS-EGFP ^dsRNA^ (BL#: 41556) and dautherless-GAL4 (da-GAL4, BL#: 27608). Balancer chromosomes from stock 27608 were substituted with autosomes from stock w^1118^. Females from the ubiquitously expressed GAL4 driver, da-GAL4, were crossed with males from both UAS-dsRNA lines. Larvae from this cross were injected and analyzed as described below.

### Insect species

We used 44 species of insects to test if they activated the melanization response. Drosophilid species (kind gift from Ben Longdon), *A. tabida* strain SFA3 (collected in Sainte Foy-Lès-Lyon, Rhône, France in 2012 and provided by Fabrice Vavre) and *L. boulardi* strain G486 [34] were lab maintained stocks. All other species were collected in Cambridge, UK, in July 2018 and identified morphologically. For large species a single specimen was collected, while for smaller species multiple individuals were pooled.

### Oil Injections

To test whether insect extracts could activate the immune response, we homogenized insects in paraffin oil. Our initial characterization of wasp extracts used 20 female *L. boulardi* in 200μl of paraffin oil (Sigma-Aldrich M5904; approximately 0.025mg wasp/μl oil). In the experiment involving multiple species, specimens were weighed and paraffin oil was added to reach a concentration of 0.025mg/μl. For large specimens the thorax was used, while for small specimens the entire animal was used (body part did not have a significant effect on melanization rates). Specimens were homogenized in the paraffin oil with a pestle in 0.5ml microcentrifuge tubes. To remove large particles, the solution was centrifuged for 1min at 300g and the supernatant was transferred to a new 0.5ml microcentrifuge tube.

Borosilicate glass 3.5” capillaries (Drummond Scientific Co. 3-000-203-G/X) were pulled to form thin needles in a needle puller (Narishige PC-10). The needle was backfilled with the oil solution with a syringe and attached to a nanoinjector (Drummond Scientific Co. Nonoject II). Late 2^nd^ instar and early 3^rd^ instar larvae were carefully removed with forceps from cornmeal food plates and placed on filter paper, in groups of 10. Larvae were carefully injected with 4.6nl of solution. After injection, ddH20 was added with a brush to remove the larvae and 40 larvae were transferred into a cornmeal food vial at 25°C, 70% relative humidity and an 8:16 dark:light cycle. After 48 hours larvae were removed with a 15% w/v sugar solution and scored for total melanization of the oil droplet.

### Hemocyte counts

To count hemocytes, larvae were injected as described above. After 48 hours, injected and control larvae were collected, washed in PBS, dried on filter paper and pooled in groups of 8 to 10 larvae in a well of a multi-well porcelain plate. Larvae were rapidly dissected with a pair of forceps from the ventral side. Hemolymph was recovered with a 1-10μl micropipette and transferred into a 0.5ml microcentrifuge tube. 1μl of hemolymph was collected, diluted in 9ul of Neutral Red solution (1.65g/L PBS – Sigma-Aldrich N2889) and thoroughly mixed. The hemolymph dilution was transferred into a counting Thoma chamber (Marienfeld #0640711) and hemocytes were counted in a total volume of 0.1μl with a 40x objective (Leica DM750). Lamellocytes were distinguished from plasmatocytes and crystal cells by morphology.

### RNA sequencing

We performed RNA-seq on flies injected with wasp homogenate or oil droplets and unchallenged flies. Hemocytes from ∼100 larvae were pooled in 100μl of PBS, 24 hours after injections. Fat body samples were dissected from 8 third instar larvae and pooled in 100ul of PBS. RNA was purified from hemolymph or fat body samples in an identical manner: 1ml TRIzol [Ambion: 15596018] was added to collected tissue and the samples were homogenized by pipetting several times. 200ul of chloroform [Fisher Scientific: C/4920/08] was added; samples were shaken for 15 seconds, incubated at room temperature for 3 minutes then centrifuged at 12,000g for 10 minutes at 4°C. The upper aqueous phase (approximately 500ul) was removed to a fresh tube and RNA was precipitated by adding 2.5 volumes of isopropanol and incubated at −20°C for 1 hour. RNA was pelleted by centrifugation, washed with 70% ethanol, and re-suspended in 15ul of nuclease free water [Ambion: AM9930]. RNA was quantified by Qubit fluorometer2.0 [ThermoFisher Scientific: Q32866] with the Qubit RNA HS Assay Kit [ThermoFisher Scientific: Q32852] and integrity was assessed by gel electrophoresis. 100-4,000ng of RNA was used for RNA-Seq library preparation.

Libraries were prepared using the KAPA Stranded mRNA-Seq Kit Illumina® platform. TrueSeq DNA Low Throughput adaptors used were from Illumina® TruSeq™ KAPA Si adaptor kit KK8701 and adaptor concentrations and the number of PCR cycles used to amplify the final libraries were adjusted to the total amount of RNA used for each library. Seven hemocyte libraries that gave a low final concentration (<2ng/ul) were re-amplified for four more cycles. Quality control of the libraries to ensure no adapter dimers were present was carried out by examining 1ul of a 1:5 dilution on a High Sensitivity DNA chip (Agilent Technologies: 5067-4626) on an Agilent 2100 Bioanalyzer. The average library size including adapters was 350bp. Sequencing was carried out at the Cancer Research UK Cambridge Institute in June 2019. All 24 libraries were multiplexed and sequenced on one lane of HiSeq4000 using 50bp single end reads.

### Differential expression tests

Sequenced RNA-seq reads were trimmed and aligned to the *D. melanogaster* genome and reads counts per gene were estimated. Using Trimmomatic v.0.36 [35], we clipped adaptors sequences, removed the first three and last three bases, filtered strings of low-quality bases found in 4bp sliding windows where quality dropped below 20 and ensured that the remaining reads had a minimum size of 36bp. We mapped the resulting reads using STAR v2.6 [36] to the *D. melanogaster* reference (r6.28) [37] attained from Flybase (FB2019_02) [38]. We prepared the genome for STAR mapping using a sjdbOverhang of 49. Then, we mapped reads using the basic option for the twopassMode parameter, filtered multi-mapped reads and sorted the remainder by coordinates. We used featureCounts [39] to compute read counts for genes using their Flybase IDs. We only considered reads with a minimum quality score of 10.

We performed differential expression tests for the fat body and hemocyte libraries separately using edgeR v.3.24.3 [40,41] and limma v.3.38.3 [42]. We only kept genes that had CPM greater than or equal to 2 in at least four samples for a given tissue. We normalized read counts using trimmed mean of M-values. For a given tissue, we had four replicate libraries for each of three groups: wasp homogenate, oil and unchallenged. Salivary gland and male germ tissues were difficult to exclude completely when dissecting larvae and isolating the fat body of *D. melanogaster*. To minimize noise in our differential expression tests attributable to this limitation, we excluded genes that had enriched expression in the aforementioned tissues. We obtained tissue level RNA-seq expression data from FlyAtlas2 [43] and calculate the tissue specificity index (Tau) [44] for each gene in the larvae and adult males separately. We then excluded tissue-specific fat body expressed genes (Tau >0.8) with greatest expression either in the larval salivary gland or adult male testes. We only excluded genes that had FPKM >1 in either of those two tissues in FlyAltas2. We fit a linear model using limma contrasting gene expression among the three groups (full model in script). After checking that the mean-variance trends followed the expected dispersion, we fit contrasts and used the *eBayes* function to uncover genes with evidence of significant differential expression between the wasp homogenate and unchallenged comparison and the oil and unchallenged comparison separately. We used the *heatscatter* function from the LSD v.4 R package to compare the log_2_FC in expression between the two comparisons. We then extracted genes with *P*-values of less than 0.05 after a false discovery rate (FDR) correction. For a few of these genes, we divided counts per million reads (CPM) for each library by the overall total across all libraries per gene to compare expression levels across samples. We plotted a heatmap of relative gene expression using the pheatmap v.1.0.12 R package. Serine proteases were named according to [45]. Log_2_CPM counts for all genes (unfiltered) in hemocyte and fat body tissues is accessible from the following gene expression browser https://arunkuma.shinyapps.io/waspapp/ (last accessed June 2022). We performed gene ontology (Huang et al., 2009) enrichment analyses on the differentially expressed genes using Flymine [46]. The genes detected in each tissue was used as the background list. Non-redundant gene ontology terms were identified using REVIGO [47] keeping FDR *P*-values < 0.05 and similarity = 0.4.

### Bulk RNA-seq deconvolution

We used the digital cytometric method CIBERSORTx [27] to infer the proportion of hemocyte clusters, which were identified in [26], in the bulk RNA-seq data. We first created a signature matrix using read counts from 2,000 highly variable genes in the scRNA-seq. Between 300 to 500 genes were used for barcoding cell types and a q-value of 0.01 was used to test for the significance of differential gene expression. Quantile normalization was disabled as recommended and a maximum conditional number of 999 was used by default. Only genes with average log_2_ expression of 0.5 were analysed. Five replicates were used to build the scRNA-seq reference file. Half of available gene expression profiles were randomly selected to generate the file. Then, we imputed cell fractions using the bulk RNA-seq read counts from hemocyte libraries with an S-mode batch correction and used 100 permutations to assess the significance of cluster inferences.

### Gene expression by qPCR

To analyze the expression by qPCR, RNA was extracted from pools of 10 larvae, 48 hours post injection. Larvae were homogenized in 250μl TRIzol [Ambion 15596018] with ∼10 1.0mm zirconia beads [Thistle Scientific] in a tissuelyser [Retsch MM300] and kept at −80°C. For RNA extraction, samples were defrosted and centrifuged for 10min at 4°C at 12,000g. 160μl of supernatant was transferred into 1.5ml microcentrifuge tubes, 62.5μl of chloroform [Fisher Scientific C/4920/08] was added, tubes were shaken for 15s and incubated for 3min. After a 10min centrifugation at 12,000g at 4°C, 66μl of the aqueous phase was transferred into a 1.5μl microcentrifuge tube, 156μl of isopropanol [Honeywell 33539] added and the solution thoroughly mixed. After 10min incubation samples were centrifuged for 10min at 12,000g at 4°C and the supernatant was removed. RNA was washed with 250μl 70% ethanol, centrifuged for 2min at 12,000g at 4°C. Ethanol was removed, samples dried, 20μl of nuclease free water [Ambion AM9930] was added and samples incubated at 45°C for 10min. cDNA was prepared from RNA samples with GoScript reverse transcriptase (Promega) according to manufacturer instructions. cDNA was diluted 1:10. Exonic primers for *D. melanogaster* immunity genes were designed in NCBI Primer-BLAST online tool (Table S4). The gene *RpL32* was used to normalize gene expression (RpL32_qPCR_F-d: 5’-TGCTAAGCTGTCGCACAAATGG-3’; RpL_qPCR_R-h 5’-TGCGCTTGTTCGATCCGTAAC-3’; Longdon et al. 2011). Sensifast Hi-Rox SyBr kit [Bioline, BIO-92005] was used to perform the RT-qPCR on a StepOnePlus system [Applied Biosystems]. Each sample was duplicated (qPCR technical replica). The PCR cycle was 95°C for 2min followed by 40 cycles of 95°C for 5s, 60°C for 30s. For one experimental replicate, we averaged the cycle threshold (*Ct*) values of 4 biological replicates (groups of 10 larvae). The relative expression of the gene of interest (GOI) was calculated as 2^-ΔΔCt^, where ΔΔ*Ct* = (*Ct*_*GOI(Treatment)*_ – *Ct*_*RpL32(Treatment)*_) - (*Ct*_*GOI(Control)*_ – *Ct*_*RpL32(Control)*_).

### Statistical analysis

The effects of different treatments on oil droplet melanization were analyzed with a quasibinomial generalized linear model, with the ratio of melanized to non-melanized oil droplets as a response and treatment as a fixed effect. We used Tukey’s honest significant difference test to compare treatments. To test differences in lamellocyte numbers with different treatments we used a one-way ANOVA with Tukey’s test to compare treatments. We compared gene expression (2^-ΔΔCt^) using a two-tailed *t*-test, correcting *P*-values with the Bonferroni method.

We used a phylogenetic mixed model to analyze the effect of extracts of 44 insect species on oil droplet melanization. This allowed us both to reconstruct ancestral states across a phylogeny, and test whether *Drosophila* has evolved to specifically recognized parasitoid wasps after correcting for the confounding effect of the insect phylogeny. The ratio of melanized to non-melanized oil droplets was the binomial response variable. Whether or not the insect was a parasitoid was treated as a fixed effect. The phylogeny was treated as a random effect, which allows the correlation between two species to be inversely proportional to the time since those species shared a common ancestor (following a Brownian model of evolution). A residual variance allowed for differences between species that are unrelated to the phylogeny. We used the phylogeny of the 44 insect species available through TimeTree (Kumar et al. 2017). The model was fitted using a Bayesian approach using MCMCglmm (Hadfield 2010) using an inverse gamma prior. We ran 10^6^ burn-in iterations followed 10^7^ iterations, sampling every 10^4^ iterations.

R v3.6/4 [49] and RStudio v1.2.5042 [50] were widely used for generating figures.

## Author’s contributions

F.J. and A.B.L. conceived the study. N.H., A.D., J.P.D. and A.B.L. collected data. M.P.H collected and identified the insect species. R.A., A.B.L. and F.J. analyzed the data. All authors contributed to interpret the data and write the manuscript.

## Data availability

Raw and processed data files used to generate figures and the lists of differentially expressed genes are available in NERC EDS Environmental Information Data Centre (https://doi.org/10.5285/06ea87f3-476d-40fd-acce-e6923e786d48). Scripts to analyze data are available in the Github (DOI: 10.5281/zenodo.6684609). Paired end reads from oil and wasp injections were deposited into the NCBI Sequence Read Archive (SRA) and can be accessed with Bioproject ID PRJNA685781.

---

**Figure S1.**
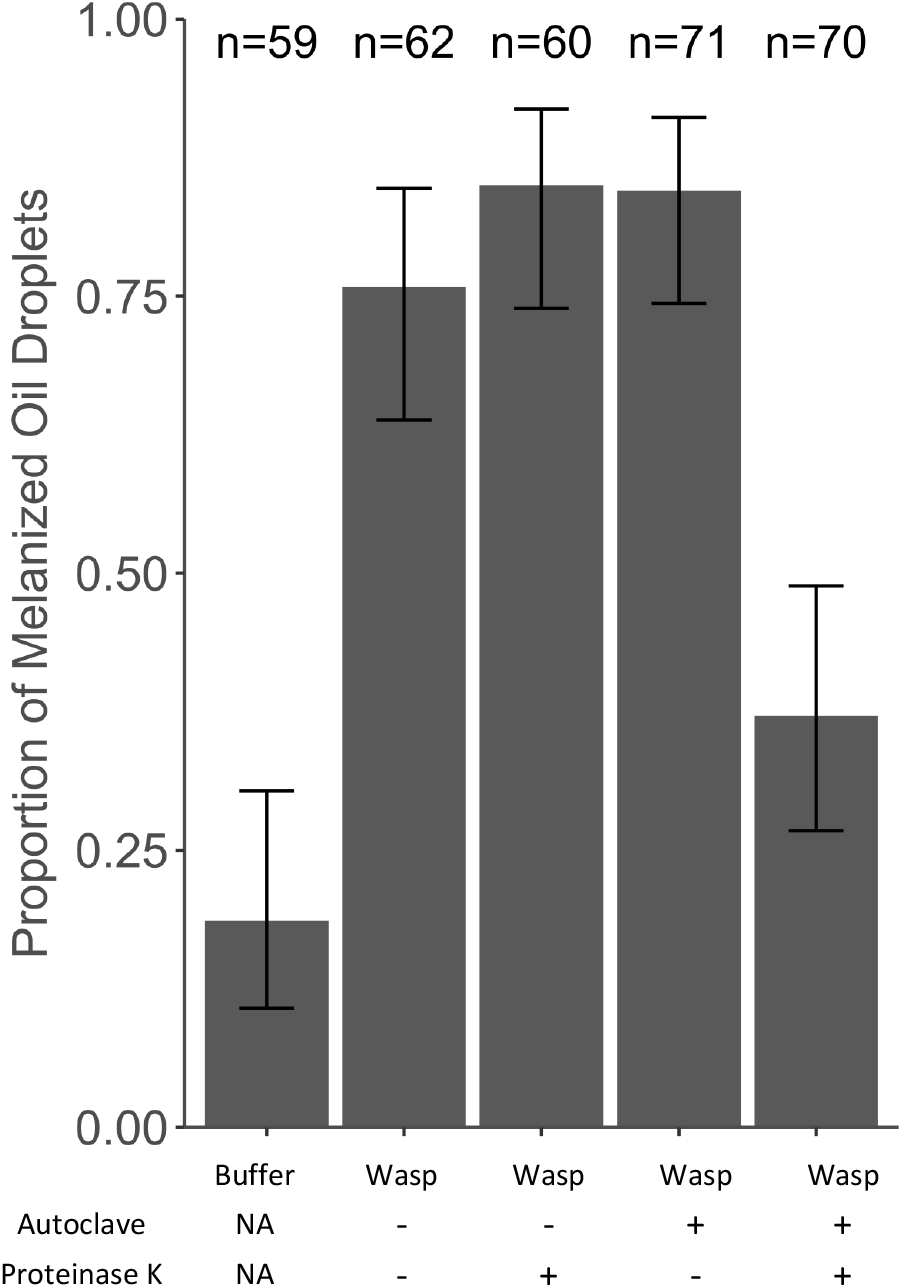
Treatment of wasp homogenate with proteinase K. Larvae were injected with oil after a first injection with buffer or wasp homogenate. Wasps were autoclaved (+) or immediately frozen (-) before homogenization in buffer. Wasp homogenates were then treated with proteinase K (+) or incubated without proteinase K (-) before injection. Number of phenotyped larvae is shown and error bars represent standard error.

**Figure S2.**
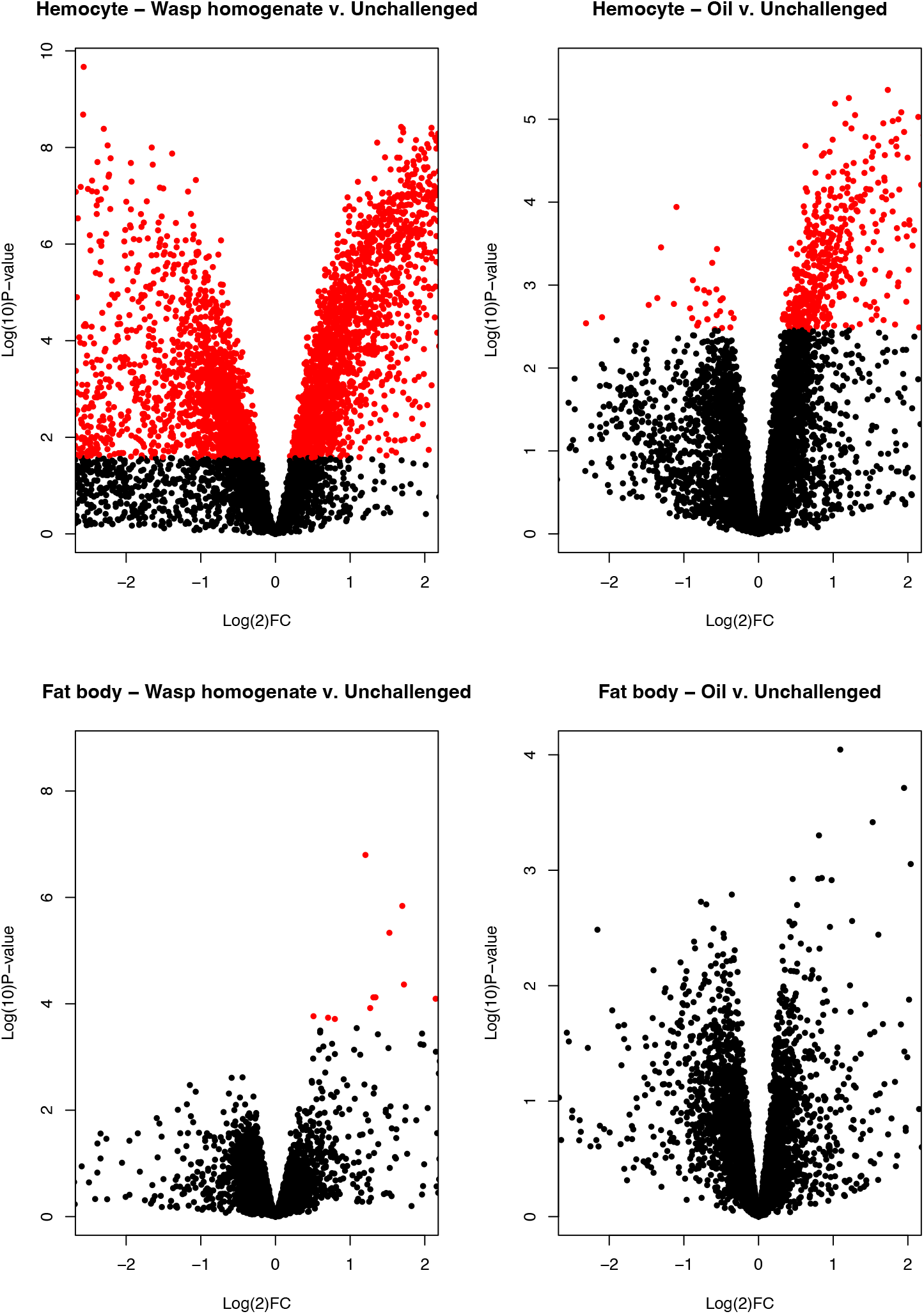
Volcano plots contrasting log_2_ fold change in gene expression against *P*-values generated from differential expression tests, for hemocyte and fat body samples. Red points indicate genes with false-discovery rate corrected *P*-values < 0.05.

**Figure S3.**
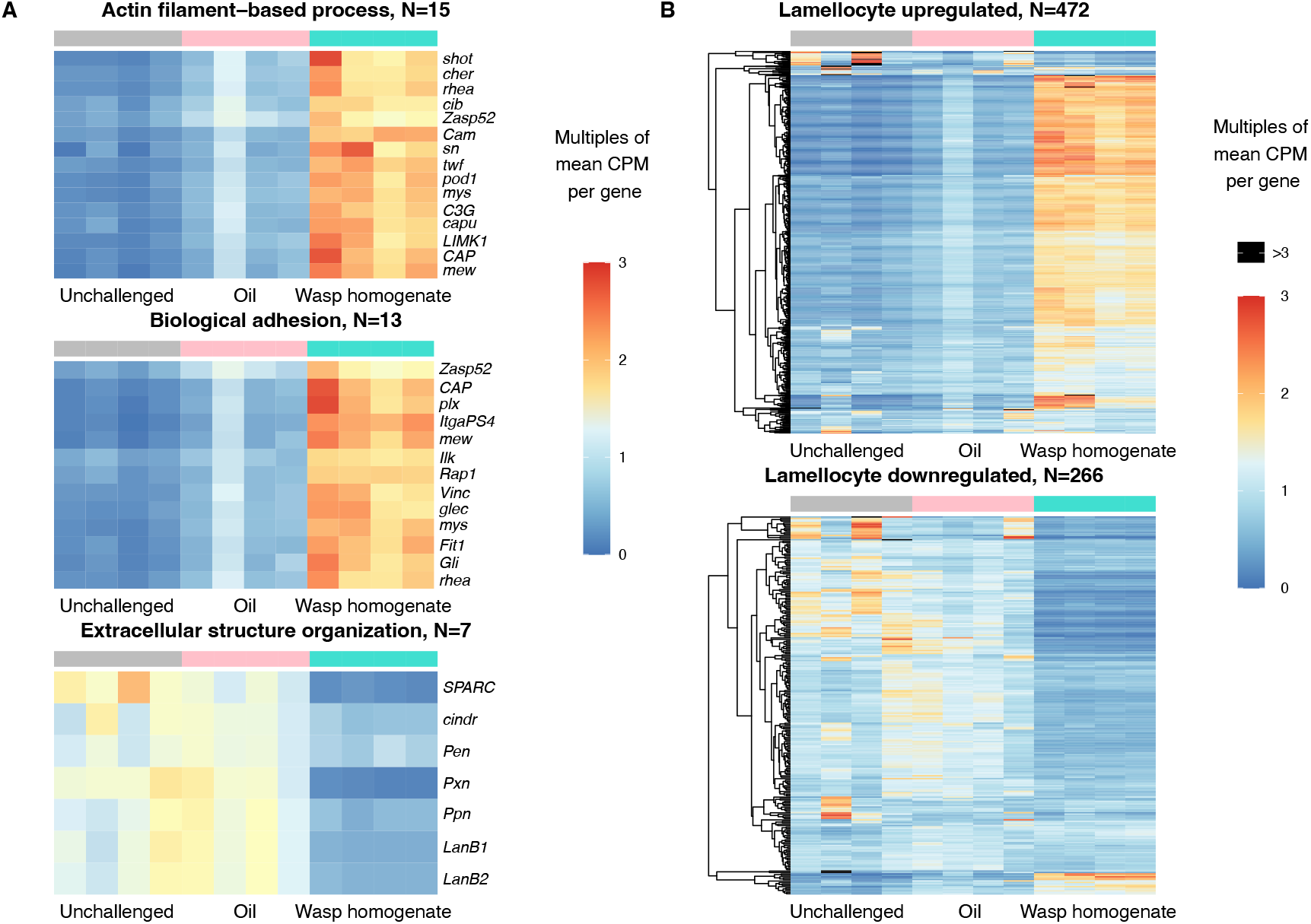
Heatmap of lamellocyte marker genes. Gene lists were attained from (Leitão et al. 2020). (**A**) Heatmap of genes part of gene ontology categories that are enriched in lamellocytes and plasmatocytes. (**B**) Heatmap of genes differentially expressed in mature lamellocytes compared to their plasmatocyte progenitors. The colour bar represents multiples of the mean counts per million (CPM) for each gene. As only few genes/samples had values >3, they are depicted as black.

**Table S1**. Quality and read mapping metrics for the 12 hemocyte and 12 fat body samples.

**Table S2**. Gene ontology (GO) terms, Kegg/Reactome pathways and Interpro protein that are significantly enriched in the fat body genes upregulated in wasp homogenate compared to unchallenged.

**Table S3**. Gene ontology (GO) terms, Kegg/Reactome pathways and Interpro protein domains that are significantly enriched in the differentially expressed genes in wasp homogenate compared to unchallenged in hemocytes. Redundant GO categories were identified using REVIGO.

**Table S4**. List of primers used for qPCR.

